# Bracken: Estimating species abundance in metagenomics data

**DOI:** 10.1101/051813

**Authors:** Jennifer Lu, Florian P Breitwieser, Peter Thielen, Steven L Salzberg

**Author notes:** To whom correspondence should be addressed Email addresses: JL FPB PT SLS.

## Abstract

We describe a new, highly accurate statistical method that computes the abundance of species in DNA sequences from a metagenomics sample. Bracken (Bayesian Reestimation of Abundance after Classification with KrakEN) uses the taxonomy labels assigned by Kraken, a highly accurate metagenomics classification algorithm, to estimate the number of reads originating from each species present in a sample. Kraken classifies reads to the best matching location in the taxonomic tree, but does not estimate abundances of species. We use the Kraken database itself to derive probabilities that describe how much sequence from each genome is shared with other genomes in the database, and combine this information with the assignments for a particular sample to estimate abundance at the species level, the genus level, or above. Combined with the Kraken classifier, Bracken produces accurate species-and genus-level abundance estimates even when a sample contains multiple near-identical species.

## Background

Metagenomics is a rapidly growing field of study, driven in part by our ability to generate enormous amounts of DNA sequence rapidly and inexpensively. Since the human genome was first published in 2001 [1, 2], sequencing technology has become approximately one million times faster and cheaper. This dramatic improvement in technology has made it possible for individual labs to generate as much sequence data as the entire Human Genome Project in just a few days.

Along with the technological advances, the number of finished and draft genomes has also grown exponentially over the past decade. There are over 4,000 complete bacterial genomes, 20,000 draft bacterial genomes, and 80,000 full or partial virus genomes in the public GenBank archive [3]. This rich resource of sequenced genomes now makes it possible to sequence uncultured, unprocessed microbial DNA from almost any environment, ranging from soil to the deep ocean to the human body, and use computational sequence comparisons to identify many of the formerly hidden species in these environments.

When it was first published in 2014, the Kraken metagenomics classifier represented a major enhancement in the speed with which large metagenomics sequence data could be processed [4], running over 900 times faster than MegaBlast [5], the closest competitor at the time. Kraken’s success and accuracy rely on its use of a very large, efficient index of short sequences of length *k*, which it builds into a specialized database. If *k* is chosen appropriately, then most sequences of length *k* in the database will be unique to a single species, and many will also be unique to a particular strain or genome. Larger values of *k* will yield a database in which even more of each genome is uniquely covered by *k*-mers; obviously, though, *k* should not be longer than the length of a sequencing read, and metagenomics projects currently generate reads as short as 75-100 base pairs (bp). Furthermore, *k*-mers must be error-free in order for Kraken to identify them, because it relies on exact matches to its database. Longer *k*-mers are more likely to contain errors, meaning that more reads will be left unclassified if *k* is too long. Smaller *k*-mers, in contrast, will yield higher sensitivity because the minimum match length is shorter. The default value of *k*=31 used by Kraken represents a good tradeoff between sensitivity and precision for current metagenomics projects.

When used to identify the taxonomic label of metagenomics sequences, the Kraken system for classification of metagenomics sequences is extremely fast and accurate [4]. When classifying raw sequence reads, though, many reads correspond to regions of two or more genomes that are identical. (The number of such ambiguous reads decreases as reads get longer.) Kraken solves this problem by assigning labels corresponding to the lowest common ancestor (LCA) of all species that share the sequence; e.g., a read might be assigned to the genus Mycobacterium, rather than to a species or a strain, because two or more species of Mycobacterium contain identical sequences that match the read.

### Ambiguity among microbial species and strains

As the database of bacterial genomes has grown, an increasing number of genomes share large portions of their sequence with other genomes. In many cases, these genomes are nearly identical; indeed, sequencing has revealed to scientists that many formerly distinct species and genuses are far closer than were known prior to sequencing. Many species have been renamed as a result, in a process that is continual and ongoing, but many other species have retained their old names, often for historical or other reasons.

For example, the species *Mycobacterium bovis* is over 99.95% identical to *Mycobacterium tuberculosis* [6], and many cases of human tuberculosis are caused by *M. bovis* (which also infects cows) rather than *M. tuberculosis* [7]. Their high sequence identity indicates that they should be considered as two strains of a single species, but they retain different species names. As a compromise, taxonomists have created the category *Mycobacterium tuberculosis complex* [8] to represent a collection of taxa that now includes more than 100 strains of five different species. This category sits above the species level but below the genus level in the current microbial taxonomy, but it can best be described as a species.

Other examples are numerous and still growing. The three species *Bacillus anthracis* (the causative agent of anthrax), *Bacillus cereus*, and *Bacillus thuringiensis* are well over 99% identical and should all be designated as a single species [9], although their names have not been changed despite their near-identity revealed by sequencing. As a compromise, taxonomists created the category *Bacillus cereus group*, between the level of species and genus, to include these three species and at least five others [10], all of which are extremely similar to one another. In some cases, two organisms that should be called the same species may even have different genus names. For example, *Escherichia coli* and *Shigella flexneri* are classified in different genera, but we know from sequence analysis that they represent the same species [11].

Failure to recognize the mutability of the bacterial taxonomy can lead to erroneous conclusions about the performance of metagenomic classifiers. For example, one recent study [12] created a mock community of 11 species, one of which was *Anabaena variabilis* ATCC 29413, not realizing that this genome had been renamed and was synonymous with species in the genus Nostoc [13]. When Anabaena was removed from the database, Kraken correctly identified the reads as Nostoc, but Peabody *et al.* erroneously considered all these reads to be misclassified.

### Classification versus abundance estimation

Kraken assigns a taxonomy label to every read in a metagenomics sample using a custom-built database that may contain any species the user chooses. Among the current set of finished bacterial and archaeal genomes, hundreds of species can be found for which large fractions of their sequence are identical to other genomes belonging to distinct strains, species, or even genera. The reads collected from these species cannot be assigned to a specific strain, and the correct behavior for a read-level classifier like Kraken is to assign them to a taxonomic level representing the lowest common ancestor (LCA) among the genomes that share the identical sequence. This implies that for some species, the majority of reads might be classified at a higher level of the taxonomy. Kraken thus leaves many reads “stranded” above the species level, meaning that the number of reads classified directly to a species may be far lower than the actual number present.

Some recent studies have erroneously assumed that Kraken’s raw read assignments can be directly translated into species-or strain-level abundance estimates [14]. This over-simplistic assumption is flawed, though, for the reasons explained here. Directly converting all read assignments to abundance estimates while ignoring reads at higher levels of the taxonomy will grossly underestimate some species, and create the erroneous impression that Kraken’s assignments themselves were incorrect.

Nonetheless, metagenomics analysis often involves estimating the abundance of the species in a particular sample. Even if we cannot unambiguously assign each read to a species, we would like to estimate how much of each species is present, either in terms of the number of reads or as a percentage of the sample. The purpose of the present study is to describe a method that combines inter-species similarity and Kraken classifications to produce species-level estimates for all species in the sample.

## Computational Methods

Our new method, Bracken (Bayesian Reestimation of Abundance of Species and Sequences), estimates species abundances in metagenomics samples by probabilistically re-distributing reads in the taxonomic tree. Reads assigned to nodes above the species level are distributed down to the species nodes, while reads assigned at the strain level are re-distributed upward to their parent species. For example, in **Figure 1** we would distribute reads assigned to family F and genus G1 down to species S1and S2, and reads assigned to strains S1_1_ and S1_2_ would be assigned to species S1. As we show below, Bracken can easily reestimate abundances at other taxonomic levels (e.g., genus or phylum) using the same algorithm.

**Figure 1.**
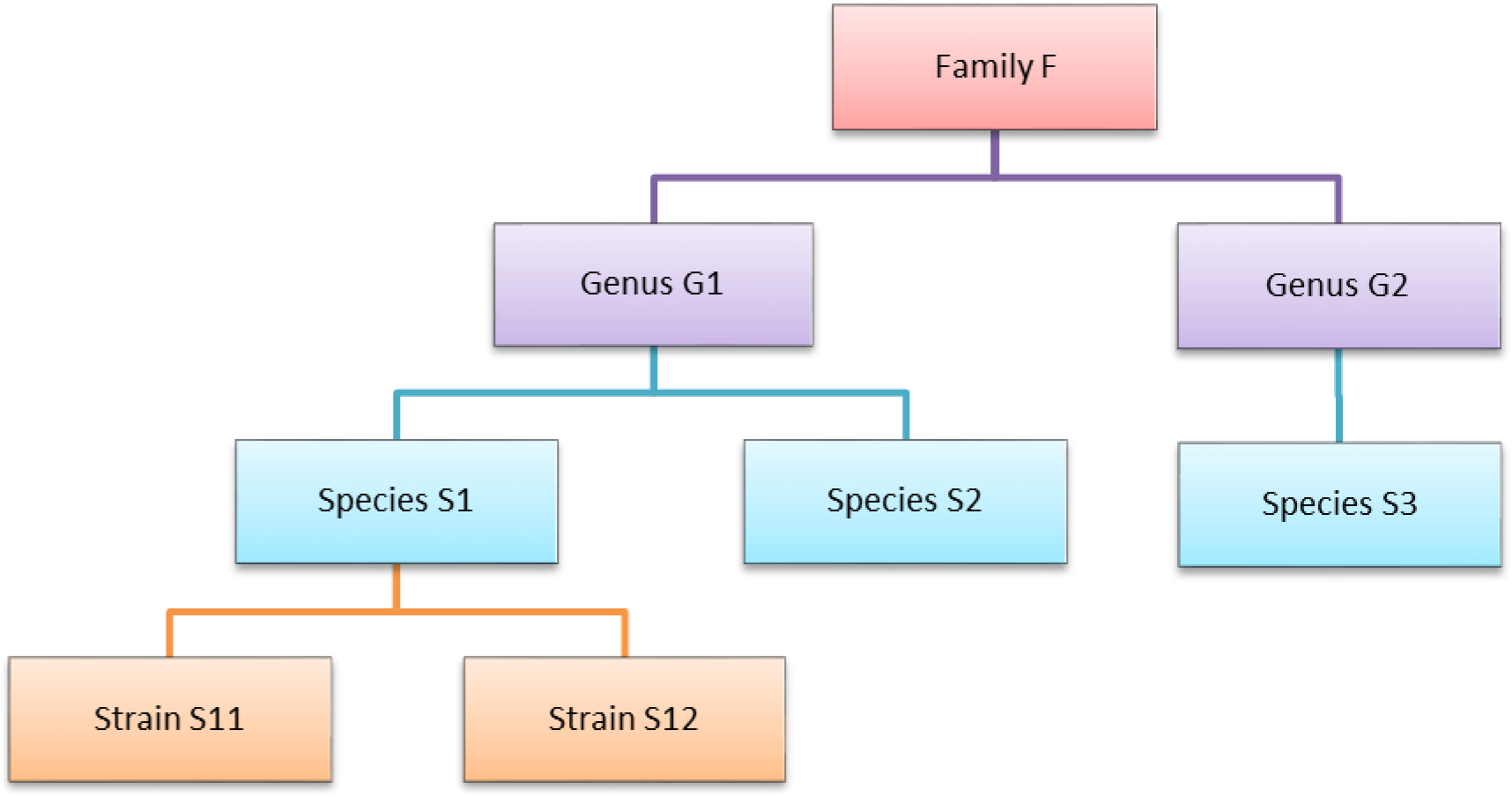
Schematic showing a small portion of a hypothetical taxonomy of genomes.

In order to re-assign reads classified at higher-level nodes in the taxonomy, we need to compute a probabilistic estimate of the number of reads that should be distributed to the species below that node. To illustrate using the nodes in **Figure 1**, we need to allocate all reads assigned to genus G1 to species S1 and S2 below it, and reads assigned to the family F would have to be allocated to species S1, S2, and S3.

Reallocating reads from a genus-level node in the taxonomy to each genome below it can be accomplished using Bayes’ theorem, if the appropriate probabilities can be computed. Let *P(S*_*i*_) be the probability that a read is classified by Kraken as genome Si, *P(G*_*j*_) be the probability that a read is classified by Kraken at the genus level G_j_, and *P(G*_*j*_ | *S*_*i*_) be the probability that a read from genome Si is classified by Kraken as the parent genus Gj.

Then the probability that a read classified at genus G_j_ belongs to the genome S_i_ can be expressed as Equation 1:

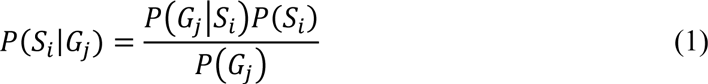

Note that because we began by assuming that a read was classified at node G_j_, *P(G*_*j*_) = 1.

Next we consider how to compute *P(G*_*j*_ | *S*_*i*_), the probability that a read from genome S_i_ will be classified by Kraken at the parent genus G_j_. We estimate this probability for reads of length *r* by classifying the sequences (genomes) that we used to build the database using that same database, as follows. For each *k*-mer in the sequences, Kraken assigns it a taxonomy ID by a fast lookup in its database. To assign a taxonomy ID for a read of length *r*, Kraken examines all *k*-mer classifications in that read. For example, for *k*=31 and *r*=75, the read will contain 45 *k*-mers. Our procedure examines, for each genome in the database, a sliding window of length *r* across the entire genome.

To find the taxonomy ID Kraken would assign to each window, we simply find the deepest taxonomy node in the set of *k*-mers in that window. Since each *k*-mer in a database sequence is assigned to a taxonomy ID somewhere along the path from the genome’s taxonomy ID to the root, the highest-weighted root-to-leaf path (and thus the Kraken classification) corresponds to the deepest node.

For each genome S_i_ of length *L*_*i*_ we thus generate *(L*_*i*_-*r+1)* mappings to taxonomical IDs. For each node G_j_, we then count the number of reads from S_i_ that are assigned to it, *N*_*Gj(Si)*_. *P(G*_*j*_ | *S*_*i*_) is then the proportion of reads from S_i_ that were assigned to the genus node G_j_; i.e., *P(G*_*j*_ | *S*_*i*_) = *N*_*Gj(Si)*_*/(L*_*i*_-*r+1)*.

The final term that we must calculate from Equation 1 is *P(S*_*i*_), the probability that genome S_i_ exists in the sample, which is computed in relation to other genomes from the same genus. For example, if the sample contains three genomes in the same genus, and if S_i_ comprises 30% of the reads from those three genomes, then *P(S*_*i*_) = 0.3. We estimate this probability using the reads that are uniquely assigned by Kraken to genome S_i_, as follows.

If we let *U*_*si*_ be the proportion of genome S_i_ that is unique, then

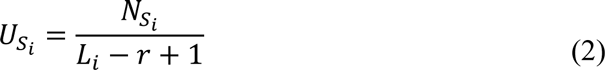

where *N*_*Si*_ is the number of k-mers of length *r* that are uniquely assigned to genome S_i_ by Kraken, and *L*_*i*_ is the genome length. For example, if *L*_*i*_ = 1 Mbp and only 250,000 k-mers are unique to genome S_i_, then *U*_*si*_ = 0.25.

Then, using the number of reads *K*_*Si*_ from a sample that Kraken actually assigns to S_i_, we can estimate the number of reads that likely derive from S_i_ as:

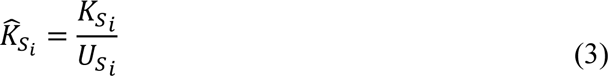

For example, if Kraken classifies 1000 reads as genome S_i_ and 25% of the reads from S_i_ are unique, then we would estimate that 4000 reads (1000 / 0.25) from S_i_ are contained in the sample.

If genus G_j_ contains *n* genomes, we estimate the number of reads 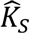 for each of the *n* genomes and then calculate *P(S*_*i*_) by:

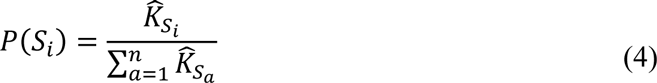

Using this result in Equation 1 above allows us to compute *P*(*S*_*i*_|*G*_*j*_) for each genome S_i_. Each probability *P*(*S*_*i*_|*G*_*j*_) is then used to estimate the proportion of the reads assigned to genus G_j_ that belong to each of the genomes below it.

These calculations are repeated for each taxonomic level above the genus level (family, class, etc.), with read distribution at each level going to all genomes classified within that taxonomic subtree.

To compute species abundance, any genome-level (strain-level) reads are simply added together at the species level. In cases where only one genome exists for a given species, we simply add the reads distributed downward from the genus level (and above) to the reads already assigned by Kraken to the species level. In cases where multiple genomes exist for a given species, the reads distributed to each genome are combined and added to the Kraken-assigned species level reads. The added reads give the final species-level abundance estimates.

This method can also estimate abundance for other taxonomic levels. In such cases, only higher nodes within the taxonomy tree undergo read distribution. After distributing reads downward, we estimate abundance for a node at the level specified by combining the distributed reads across all genomes within that node’s subtree.

## Results and Discussion

We applied the statistical re-assignment method described here to create species-level abundance estimates for several metagenomics data sets. The overall procedure works as follows. First, we compute a set of probabilities from the Kraken database by computing, for every sequence of length R in every genome, where it will be assigned in the taxonomy (see Methods). For our experiments we set R=75 as our datasets contain 75-bp reads. Bracken can use these probabilities for any metagenomics data set, including data with different read lengths, although the estimates might be slightly improved by recomputing with a read length that matches the experimental data.

Second, we run Kraken on the dataset to produce read-level taxonomic classifications. We then apply our abundance estimator, Bracken, which uses the numbers of reads assigned by Kraken at every level of the taxonomy to estimate the abundances at a single level (e.g., species). Note that to exclude false positives, Bracken ignores species with counts below a user-adjustable threshold.

### Experiments on a 100-genome metagenomics data set

For our first experiments, we used a data set containing simulated Illumina reads from 100 genomes. This data, which we call here the ilOO dataset, was used previously in a comparison of metagenomic assembly algorithms [15]. The data contains 53.3 million paired reads (26.7M pairs) from 100 genomes representing 85 species. The reads have error profiles based on quality values found in real Illumina reads [15]. The ilOO dataset includes several very challenging genomes for this task, including multiple strains and species in the genuses Bacillus and Mycobacteria, some of which are nearly identical to one another. The ilOO data are freely available at http://www.bork.embl.de/~mende/simulated_data.

The difficulty of estimating species abundance increases as the database itself contains more species. For example, it would clearly be easier to estimate abundances in the i100 dataset if we used a Kraken database containing only the 100 genomes in that dataset. To make the problem more realistic, we built two different databases and estimated abundance using both. The first (“small”) database contains 693 genomes including the i100 genomes; this is the full database from the simulation study by Mende *et al.* [15]. The results when using the small database for classification are shown in **Figure 2**. For several species, the initial Kraken numbers (reads assigned to a particular species) are far too low, because many of the reads (for some genomes, a large majority) were assigned labels at the genus level or above. After reestimation with Bracken, these reads were redistributed to the species level, with the result that almost all the abundance estimates were 98-99% correct, as shown in the figure.

**Figure 2:**
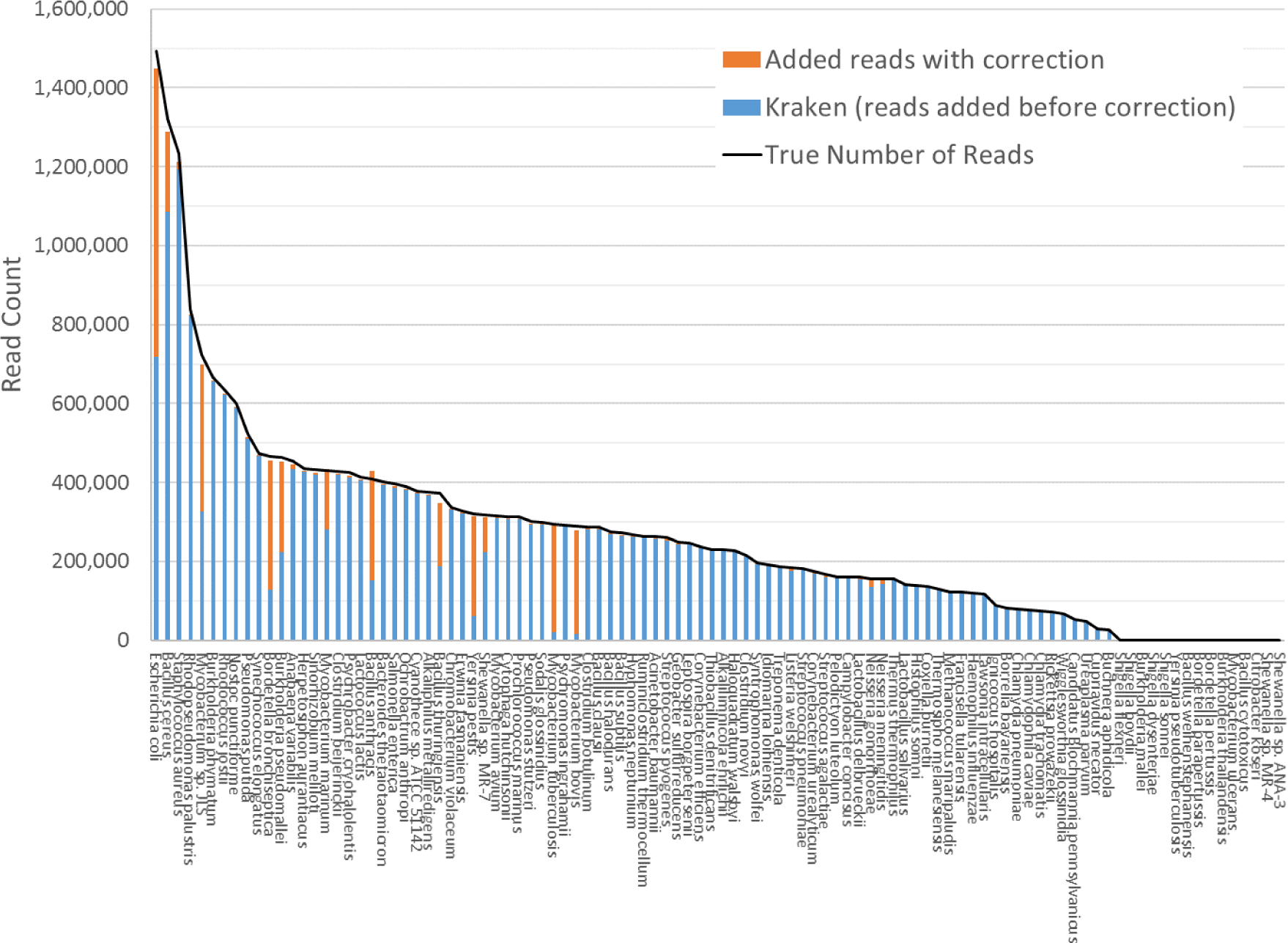
Estimates of species abundance in the ilOO metagenomics dataset computed by Kraken (blue) and Bracken (blue+orange). For this result, the Kraken database contained 693 genomes that included the ilOO genomes. The black line shows the true number of reads from each species.

The second (“large”) database contains all genomes used in the synthetic and spike-in experiments, as well as a broad background of bacterial genomes. In particular, it includes all complete bacterial and archaeal genomes from RefSeq as of 25 July 2014 (archived at ftp://ftp.ncbi.nlm.nih.gov/genomes/archive/old_refseq), which total 2596 distinct taxa, plus those i100 genomes that were not present in the RefSeq data. We also added the nine genomes used in our skin bacteria spike-in experiment (described below) resulting in a total of 2635 distinct taxa. The complete list of sequences in the large database can be found in Supplementary Table 1. The resulting Kraken database has a size of 74 gigabytes.

Figure 3 shows results when using the large database to estimate abundance for the i100 genomes. This test is much more difficult because of the large number of similar and near-identical genomes in the database. Many more reads are ambiguous, mapping identically to two or more species, which means that Kraken assigns them to the LCA of those species. Nonetheless, Bracken brings the estimated abundance of all species within 4% of the true abundance, and most fall within 1%. Supplementary Tables 2A-B contains the detailed numbers for all species in **Figures 1** and **2**.

**Figure 3:**
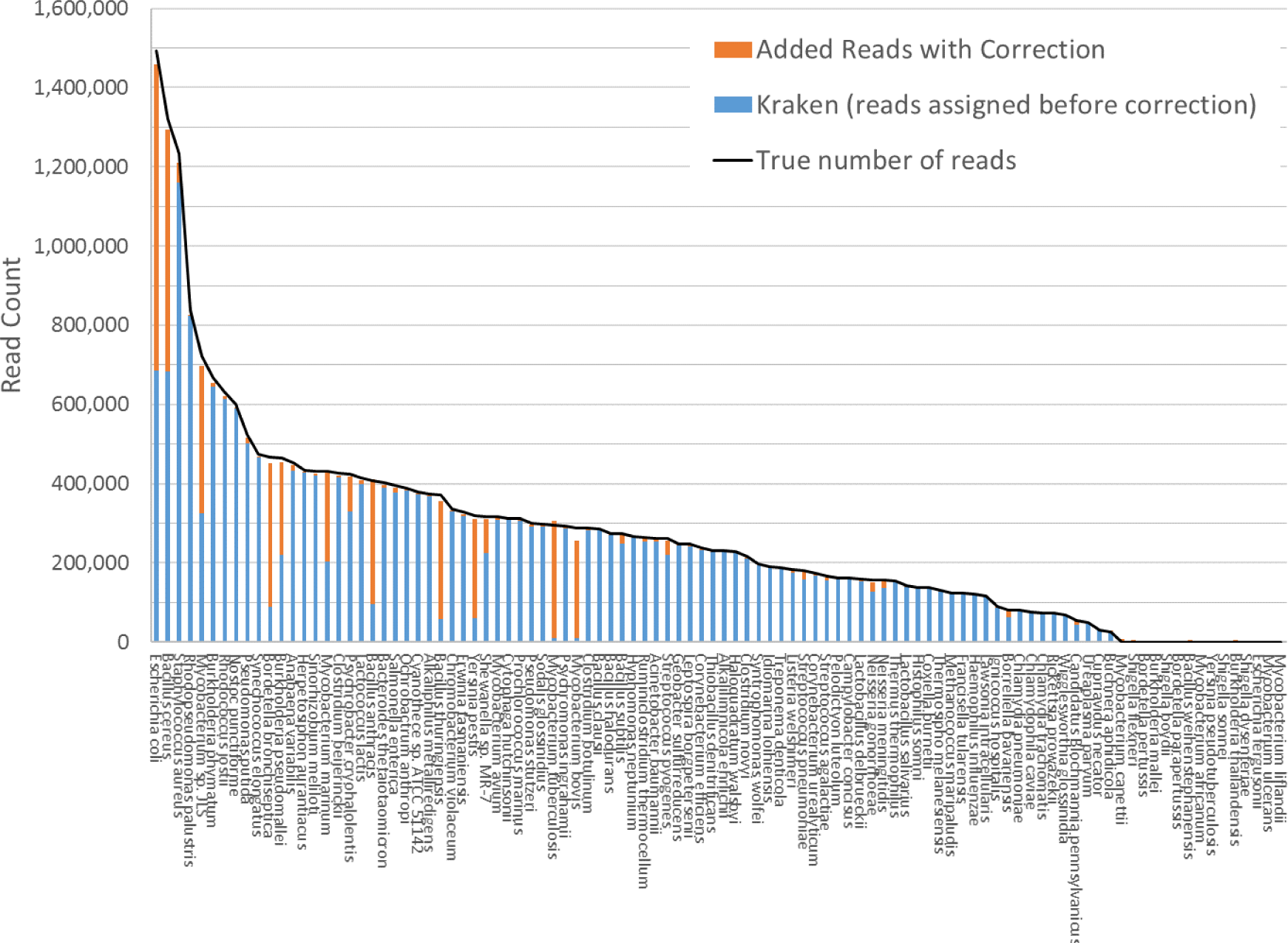
Estimates of species abundance computed by Kraken (blue) and Bracken (blue+orange) for the i100 metagenomics data. For this result, the Kraken database contained 2635 distinct bacterial and archaeal taxa. The black line shows the true number of reads from each species.

Within the i100 genomes, the five species belonging to the Mycobacterium genus (*M. tuberculosis, M. bovis, M. avium, M. marinum*, and *M. sp. JLS*) pose a particular challenge for abundance estimation due to the similarities among their individual genomes. For example, Kraken classified only 9,733 *M. tuberculosis* reads at the species level, and classified the remaining 285,414 reads as either Mycobacterium (a genus) or *M. tuberculosis complex* (a taxonomic class intermediate between genus and species), as shown in **Figure 4** and Supplementary Table 3. For these Mycobacteria genomes, Bracken reallocated the reads from higher-level nodes to yield species abundance estimates within 4% of the true abundance. **Figure 4** shows the number of reads assigned to each species by Kraken, the true number of reads, and the number of reads assigned to each species by Bracken after abundance reestimation. As shown in Supplementary Table 3, for *M. tuberculosis* Bracken redistributes the reads to produce an estimate of 306,792 reads, close to the true value of 295,147.

**Figure 4.**
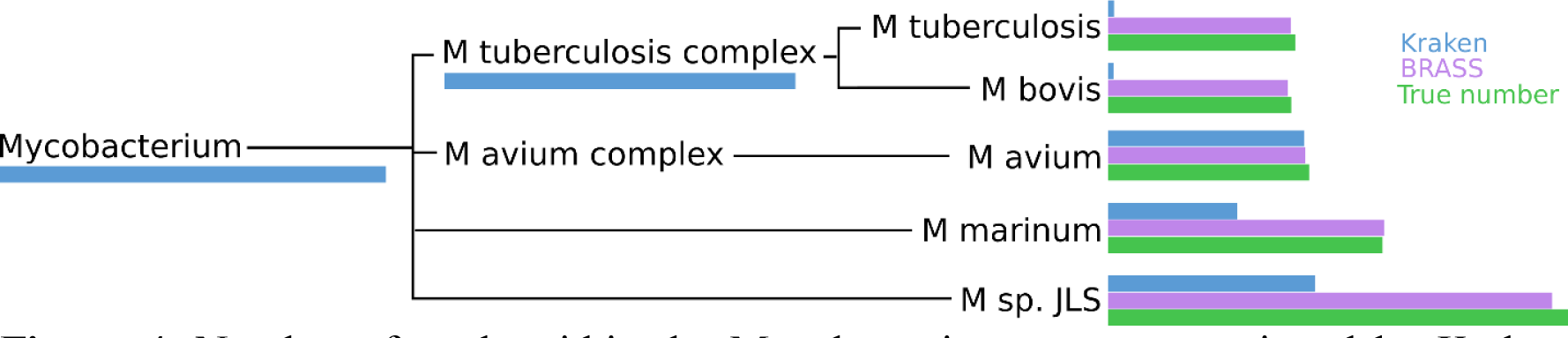
Number of reads within the Mycobacterium genus as assigned by Kraken (blue), estimated by Bracken (purple) and compared to the true read counts (green).

Much of the ambiguity disappears when classifying reads at the genus level or above. Bracken can estimate abundances at any taxonomic level, from species to phylum. To illustrate, we used it to estimate the abundance of each genus in the i100 data, again using the large database. **Figure 5** shows that Bracken’s abundance estimates for all genuses are 99% accurate or better. It also shows that for all but three genuses, the initial Kraken counts alone were equally accurate. Note that a similar plot for family, class, order, or phylum would show essentially perfect accuracy, as there were very few reads that were ambiguous at these higher levels.

**Figure 5.**
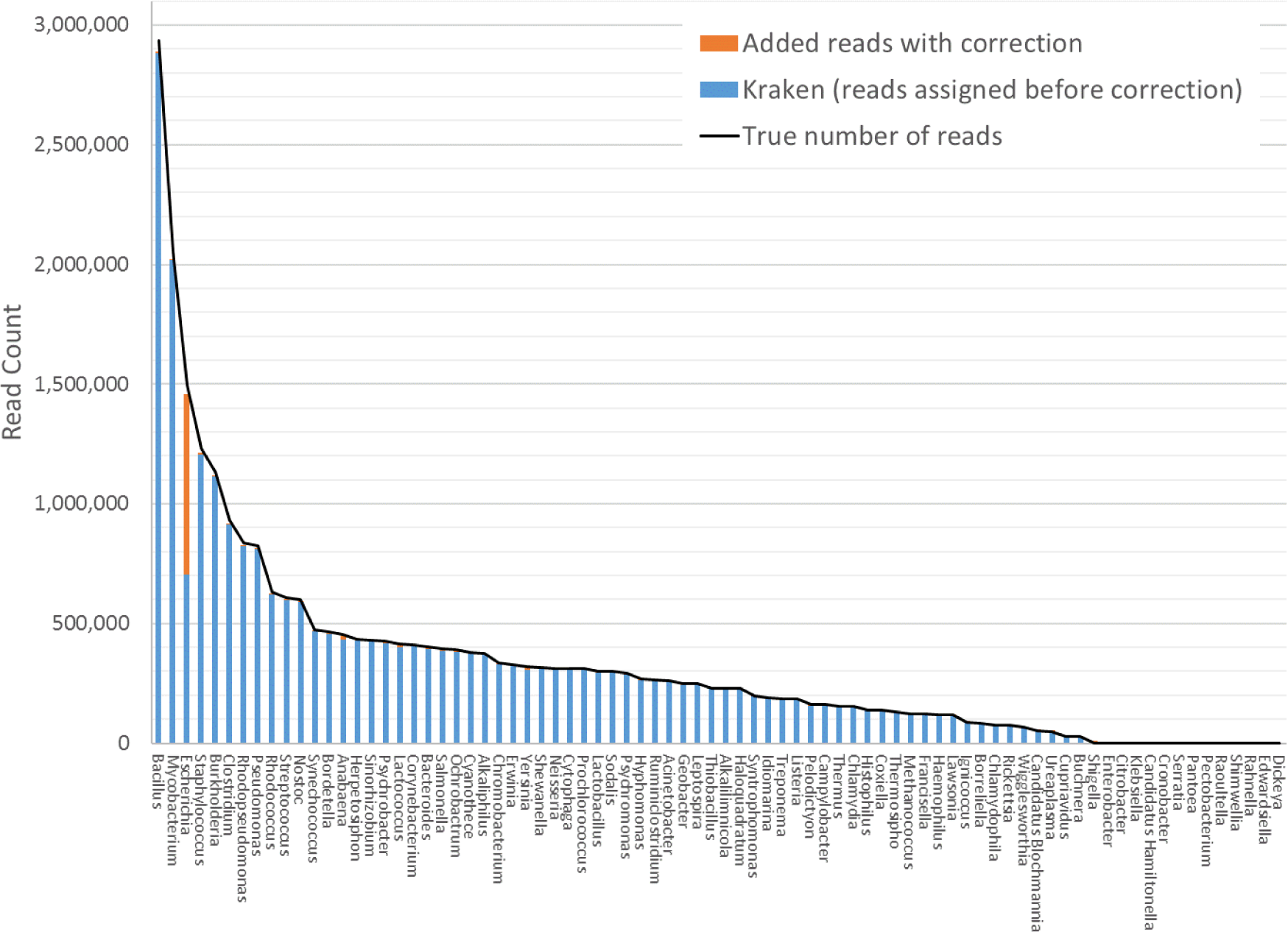
Estimates of genus abundance in the i100 metagenomics sample computed by Kraken (blue) and Bracken (blue+orange). For this result, Kraken used the large database of 2635 taxa. The black line shows the true number of reads from each genus.

### Experiments on a real metagenomics sample created from known species

For a more realistic evaluation of the performance of Bracken, we generated new sequence data using a set of bacteria that are commonly found on healthy human skin. This mock community was assembled by combining purified DNA from nine isolates that were identified and sequenced during the initial phase of the Human Microbiome Project [16]: *Acinetobacter radioresistens* strain SK82, *Corynebacterium amycolatum* strain SK46, *Micrococcus luteus* strain SK58, *Rhodococcus erythropolis* strain SK121, *Staphylococcus capitis* strain SK14, *Staphylococcus epidermidis* strain SK135, *Staphylococcus hominis* strain SK119, *Staphylococcus warneri* strain SK66, and *Propionibacterium acnes* strain SK137. We used Kraken with the large database to classify all 78.4 million read pairs from this data set (see Methods).

We used Bracken to estimate both species and genus-level abundance in the skin microbiome community. In the Bracken results, the nine true species comprise over 99% of the species-level abundance estimates. The mixture was created with approximately equal amounts of each of the nine genomes, so the expectation was that each species would account for ~11% of the total. However, as shown in **Figure 6**, the estimates varied from 7.3% to 14.8%. Details for the exact number of reads assigned by Kraken and the abundance estimates by Bracken are shown in Supplementary Table 4.

**Figure 6.**
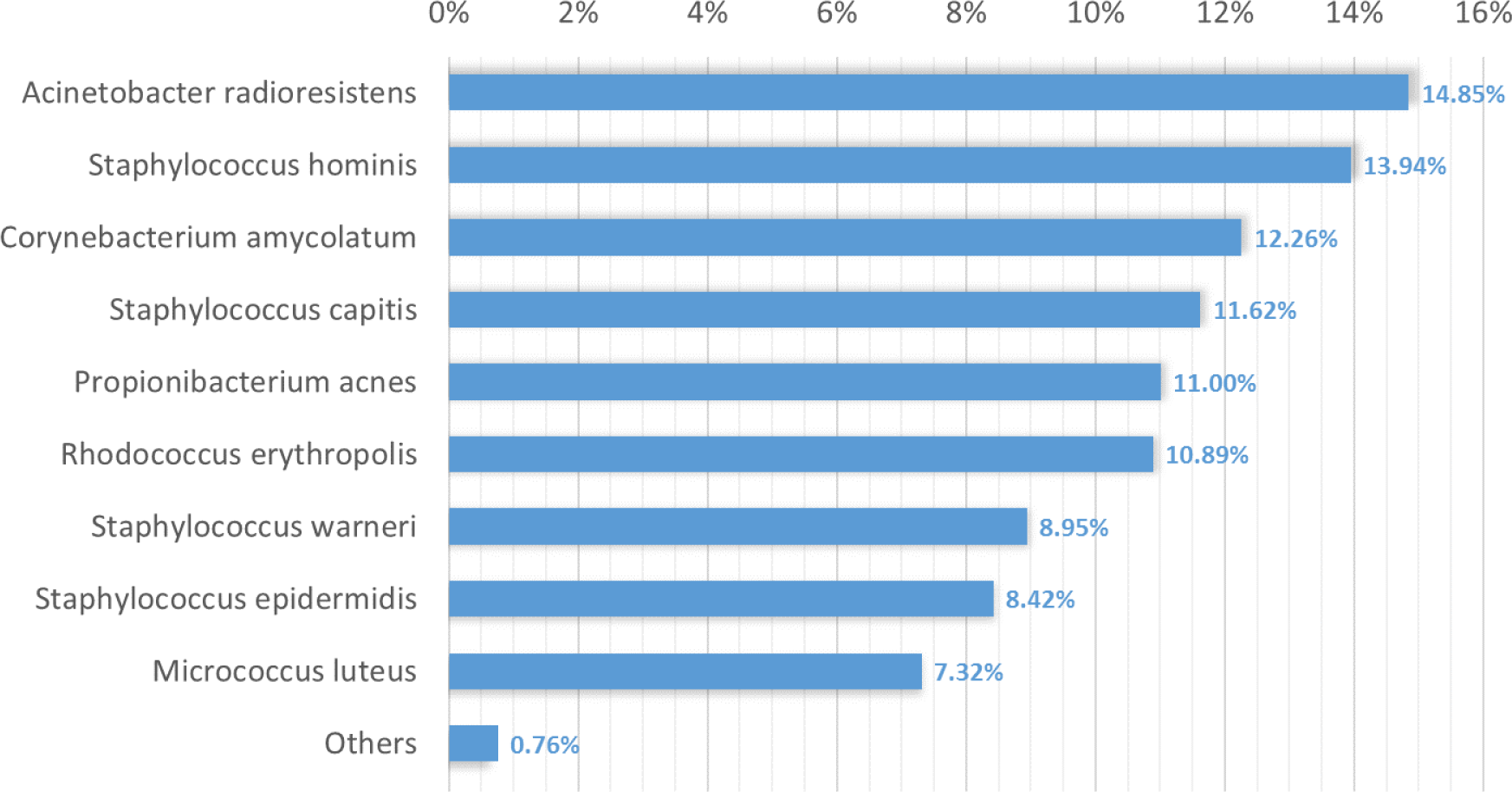
Estimates of species abundance made by Bracken for the metagenomics community containing isolates of nine bacterial species commonly found on human skin.

Deviations from the expected abundance could arise from a variety of factors. The process of quantifying DNA and mixing in equal amounts can be influenced by pipetting consistency. Second, library amplification by PCR, an integral step in the Nextera library preparation process, can exaggerate small differences in quantities and lead to significant biases in abundance [17]. We examined a sample of the classified reads by hand, and could find no evidence that Kraken mis-classified reads from M. luteus (the smallest portion of the community, estimated at 7.3%) to any of the other species or genuses. The abundances found in this data, therefore, may correspond fairly closely with the true abundances.

The genus-level abundance estimates computed by Bracken also correspond closely to the expected abundances for the six genuses included in the sample. Four of the nine species belong to the genus Staphylococcus, which was thus expected to comprise 44% (4 × 11%) of the sample. The Bracken estimate was 43.3%. Each of the other genus classifications has only one species present, and their abundance estimates are the same for both genus and species.

The comparison between the Kraken classification of reads and Bracken’s reassignment revealed that the nine species are sufficiently distinct to allow Kraken to classify a large majority of reads at the species level, with very few reads being classified at higher levels of the taxonomy. Specifically, Kraken classified 76.4 million reads to the nine species included in the sample. Only 1.3 million reads out of the 78.2 million total (1.6%) were classified by Kraken at the genus level or above. (The remaining reads were unclassified.) In this case Bracken does not provide a substantial benefit, because reassignment of the 1.3 million reads could yield at most a 1.6% change in the estimated composition of the sample.

## Conclusion

Estimating the abundance of species, genuses, phyla, or other taxonomic groups is a central step in the analysis of many metagenomics datasets. Metagenomics classifiers like Kraken provide a very fast and accurate way to label individual reads, and at higher taxonomic levels such as phyla, these assignments can be directly translated to abundance estimates. However, many reads cannot be unambiguously assigned to a single strain or species, for at least two reasons. First, many bacterial species are nearly identical, meaning that a read can match identically to two or more distinct species. Second, the bacterial taxonomy itself is undergoing constant revisions and updates, as genome sequencing reveals the need to re-assign species to new names. These revisions sometimes create new taxa that share near-identical sequence with a distinct species. In these situations, Kraken correctly assigns the read to a higher-level taxonomic category such as genus or family. This creates a problem in that Kraken’s classifications cannot be used directly for species abundance estimation.

Bracken addresses this problem by probabilistically re-assigning reads from intermediate taxonomic nodes to the species level or above. As we have shown here, these re-assignments produce species-level abundance estimates that are very accurate, typically 98% correct or higher. For genus-level abundance, accuracy is even higher because fewer reads have ambiguous assignments at that level. For abundance estimation at higher levels, ranging from family up to phylum, Kraken’s original read assignments can be used directly to create abundance estimates.

## Methods

### DNA preparation and sequencing

To generate the skin microbiome community, purified DNA was obtained from the Biodefense and Emerging Infections Research Resources Repository (BEI Resources). Each of the nine bacterial isolates was grown under conditions recommended by BEI Resources, collected by centrifugation during log growth phase at a 600nm optical density (OD^600^) of 0.8-1.2, and genomic DNA was isolated using MasterPure DNA isolation reagents (Epicentre). Purified genomic DNA was quantified using the high sensitivity picogreen assay (Invitrogen), pooled in equal amounts by mass, and prepared for sequencing using Nextera XT library preparation reagents (Illumina). The sample was then sequenced on a HiSeq sequencer, generating a total of 78,439,985 million read pairs (157 million reads), all of them 100 bp in length. These were then classified as pairs by Kraken, which concatenates the two reads from each pair and assigns them to a single taxonomic category.

### Relative Error

Supplementary Tables 2A-B include the error rate for each species in the i100 data, expressed as the difference between the true and estimated proportions. We calculated the average error as:

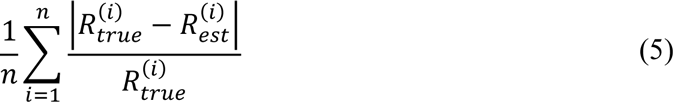

where *n* is the number of species in the i100 data, 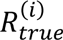 is the true number of reads for species *i*, and 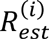 is the Bracken estimate of the number of reads for species *i*. When using the small database, the average relative error of Bracken is 1.75% across all 85 species in the i100 data. For the larger database, the average relative error is 1.89%.

## Declarations

### Software and data availability

Bracken is written in Perl and Python and is freely available for download at http://ccb.jhu.edu/software/bracken/. The reads from the skin microbiome experiment are freely available from NCBI under BioProject PRJNA316735.

## Authors’ contributions

JL and FPB wrote the software and performed the experiments. PT generated the skin microbiome HiSeq reads. JL, FPB, and SLS designed the algorithm. JL, FPB, PT, and SLS wrote the paper. All authors read and approved the final manuscript.

## Acknowledgements

Thanks to Kasper Hansen for helpful comments and feedback on a draft version of this manuscript. This work was supported in part by the U.S. National Institutes of Health under grants R01-HG006677 and R01-GM083873, and by the U.S. Army Research Office under grant W911NF-1410490.

